# A comparison of live versus kill pitfall traps to assess the diet of carabids through a metabarcoding approach

**DOI:** 10.1101/2023.03.22.532730

**Authors:** Yohann Graux, Marina Querejeta, Sabrina Gaba, Vincent Bretagnolle, Stéphane Boyer

## Abstract

Metabarcoding approaches are powerful tools to unravel trophic relationships between predators and prey. To apply metabarcoding analyses on invertebrate gut contents, specimens must be well preserved from DNA degradation, thus the trapping method should be selected accordingly. Dry pitfall traps are commonly assumed to provide a better DNA preservation than traps that use a killing agent. However, this assumption has never been specifically tested for gut content analyses.

In our study, we compared how two types of pitfall trapping, dry vs. with brine, affect the conservation of prey DNA contained in the digestive tract of predators and subsequent metabarcoding analyses. We placed dry and “classic” pitfall traps in oilseed rape fields within an intensive agricultural area in the French Nouvelle-Aquitaine region. Traps were set up in autumn and compared for carabid trapping efficiency as well as our capacity to retrieve dietary information from the digestive tract of the main carabid species, *Nebria salina* and *Calathus fuscipes*.

PCR success rate was higher in dry pitfall traps compared to classic ones for *N. salina*. We hypothesise that this was due to the presence of PCR inhibitors in the gut of this species. The ability to sequence prey DNA did not differ between specimens caught in both trap types. The list of predated species was similar between both trap types. However, sequencing yielded more prey OTUs from specimens caught in dry pitfall traps, leading to difference in prey community composition and a greater ability to reconstruct prey community

Our analyses also shed light on the prey spectrum of *Calathus fuscipes* and *Nebria salina* in oilseed rape in autumn.

## 1 Introduction

Carabid beetles are generalist predators and have been considered for years as important agents of biocontrol in agroecosystems (**Thiele, 1977; Kromp, 1999; Symondson et al., 2002**). In oilseed rape crops for instance, carabids can predate some of the major pests such as *Meligethes aeneus* (Fabricius) (Col. Nitidulidae), *Ceutorhynchus assimilis* (Paykull) (Col. Curculionidae), and *Dasineura brassicae* (Winnertz) (Dipt. Cecidomyiidae) (**Warner, 2001;** **Büchs** & **Nuss, 2000**). However, measuring accurately their efficiency as pest regulator remains elusive (**De Heij** & **Willenborg, 2020**).

To study carabid prey spectrum, large-scale temporal and spatial sampling is often conducted in order to collect a large number of specimens and allow representative coverage as well as robust statistical analyses, e.g. on diet seasonal dynamics or the impact of some environmental parameters. Pitfall traps are typically used for the sampling of carabids and other epigeal invertebrates, which allows estimating activity-density of various ground-active arthropod taxa such as ground beetles, rove beetles, spiders or ants (**Woodcock, 2005)** or species assemblage at community level (**Rainio** & **Niemalä, 2003**). Moreover, pitfall trapping is a cheap and easy-to-use method, though leading to long-lasting debate about the capture effectiveness and the potential bias that may be encountered (**Adis, 1979; Coddington et al., 1991; Woodcock, 2005; Yi et al., 2012**). Although **Hohbein** & **Conway (2018)** have proposed a standard use of pitfall traps for obtaining invertebrate abundance indices, the design of the trapping method has to be adapted to the research question and the targeted organisms. Consequently, much variation in the trap design, material, colour, size, use of funnels and rain guards, and duration have been noted (**Weeks** & **McIntyre, 1997;** **Brown** & **Matthews, 2016**). One additional point of variation, which has caused much debate, is the use of a killing agent inside the trap.

Indeed, in order to minimise the financial costs and logistic effort of sampling, various killing agents have been tried (**Hohbein** & **Conway, 2018**), i.e. formalin, picric acid or ethylene glycol. These are now rarely used due to hazardous effects on human health or wildlife (**Hall, 1991**). More recent agents, such as propylene glycol, ethanol, brine or water (at varying concentrations) were compared for their efficiency to capture rate (**Schmidt et al., 2006;** **Topping** & **Sunderland, 1992;** **Koivula et al., 2003;** **Curtis, 1980**), morphological preservation (**Braun et al., 2009; Sasakawa, 2007**; **Aristophanous, 2010**) and DNA preservation (**Vink et al., 2005;** **Gurdebeke** & **Maelfait, 2002**).

The choice of preservatives is particularly crucial when the trapped samples are to be used for subsequent genetic analysis and, most importantly, molecular sequencing. For optimal results, molecular methods require specimens in which the DNA is well-conserved. Consequently, the main criteria when selecting a sampling method for such studies is to protect the DNA from chemical and enzymatic degradation. Similarly, conservation methods used subsequently to the trapping, can also cause DNA degradation, for example the hydrolytic and oxidative effects of ethanol, especially at low concentration and high temperature (**Vink et al., 2005**). Freezing or drying insect specimens are the most suitable methods to preserve the DNA in a sample (**Post et al., 1993; Reiss et al., 1995**). However, these are difficult to apply in the field.

Potential strategies to limit the DNA degradation include using a killing agent that limits DNA degradation during the sampling period (**Pokluda et al., 2014; Stoeckle et al., 2010; Reiss et al., 1995**) and/or reducing the duration of the sampling. When the duration of the sampling is greatly reduced, it is possible to operate without preservatives and thus use dry pitfall traps, where the killing agent is replaced by a non-lethal substrate and the trapped specimens stay alive until they are picked up. Such traps are often preferred when specimens are used for metabarcoding analysis of carabid’s gut contents (e.g., **Kamenova et al., 2018a; Roubinet et al., 2017; Frei et al., 2019**), especially because they allow to obtain regurgitates, which give a higher prey DNA detection success for large amplicons (**Waldner** & **Traugott, 2012**). Because dry pitfall traps prevent the samples to be submerged in a trapping agent that is potentially deleterious for the prey DNA contained in the gut, we may assume that prey DNA inside the gut of predators is better preserve in those traps. However, by keeping predators alive, the enzymatic reactions inside their digestive tract are maintained after the trapping and this could lead to greater degradation of prey DNA. Therefore, it remains unknown whether dry traps affect our ability to retrieve robust information about prey-predator interactions.

Our aim here was to compare two types of pitfall trapping, dry vs. with brine (here called “classic”), regarding the conservation of prey DNA contained in the digestive tract of predatory species and subsequent analyses. To do this, we used a field-based approach together with a metabarcoding approach to analyse the diet of Carabids captured in 21 oilseed rape fields with either dry or classic traps, and compared the results to assess the impact of the trapping method on our capacity to retrieve dietary information from metabarcoding.

## 2 Material and methods

### 3 Carabids collection and sample processing

We captured carabid beetles from 21 oilseed rape crops located in an intensive agroecosystem within the Long-Term Social-Ecological Research *Zone Atelier* “Plaine & Val de Sèvre” (hereafter ZAPVS, Nouvelle-Aquitaine; see **Bretagnolle et al., 2018a**). Samples were collected in autumn, during two sampling periods in October 2020 (5^th^ to 9^th^ and 26^th^ to 30^th^). In each field, 10 pitfall traps (**Figure 1a**) were used, five were dry (half filled with clay beads) and five were classic pitfall traps (half filled with 200 mL of water, 10 g/L of salt and 5 drops/L of inodor soap). The classic pitfall trap used here is the main type of trap used for biodiversity monitoring in the ZAPVS (**Bretagnolle et al., 2018b**). The traps were positioned on two lines in a staggered arrangement (**Figure 1b**). Because 90% of carabid species reproducing in autumn are active at night (**Thiele** & **Weber, 1968**), the traps were activated for one night between 17 P.M. and 8 A.M. of the next day. In this way traps were in activity for a maximum of 15 hours.

**Figure 1:**
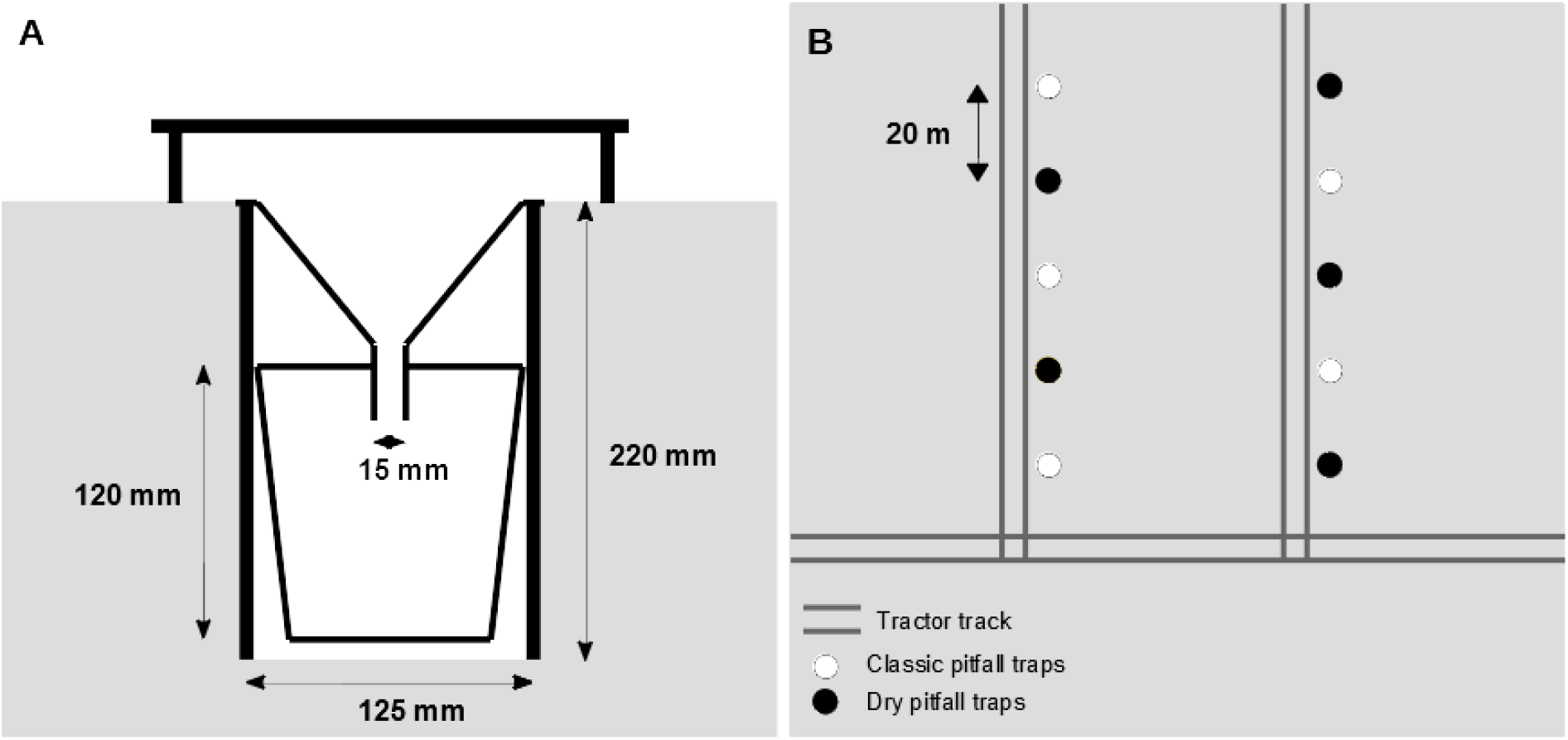
Pitfall trap design (A). Traps were filled to half the height with salted water and inodor soap (classic trap) or with clay beads (dry trap). Pitfall traps staggered arrangement in the field (B). The parallel tractor tracks are separated by approximately 40 m.

Each pitfall trap consists of a 1,000 mL plastic cup associated with a funnel (125 mm at the wide end and 15 mm at the narrow end) buried in the ground with the lip of the funnel flush with the soil surface. A plastic plate provided shade and protection from rain (**Figure 1**).

All trapped invertebrates inside the dry pitfall traps were collected in the morning, placed in a cool box for transportation and stored at −20°C within 3 hours. After 2-3 hours of freezing, carabid specimens were sorted out, identified and sexed, and placed individually in 1.5 mL tubes filled with 300 mL of 100% ethanol. Carabid specimens trapped in classic pitfall traps were first sorted, quickly dried on absorbent paper, identified and sexed, and also placed individually in 1.5 mL tubes with 300 mL of 100% ethanol. All samples were stored at −20°C until subsequent processing.

Out of 611 captured specimens, 292 were used for the gut contents extraction (162 trapped with classic pitfall traps and 130 with dry pitfall traps). Selection was based on molecular diet analyses goals, that was only conducted on specimens belonging to the dominant carabid species (i.e. those representing >10% of trapped individuals). In addition, to maximise representativeness of the different traps and the different fields, no more than 8 individuals per trap and 30 individuals per field were kept for subsequent analyses. Gut contents were isolated by the dissection of the crop, a bulb-like organ situated between the oesophagus and the proventriculus. This part of the foregut is particularly suitable for the study of the diet because it is a mechanically functioning unit ensuring filtration and food storage (**Jaspar-Versali *et al*., 1987**) and because it is made of a thick epithelium, which makes it more easily extractable than the other parts of the digestive tract. Dissections were performed with sterilised forceps. Between each sample, forceps were successively dipped in bleach, Decon™ 2%, MicroBeads sterilizer (300°C – Fisher) and then sprayed with ethanol 96%. This sterilisation process lasted about 45 minutes before the same forceps was used again for another sample. Two forceps were used for the dissection and a third one was exclusively used to harvest the crop without touching the outer surface of the beetle.

The dissected crop was emptied in 300 *μ*L of lysis buffer from the NucleoMag Tissue kit for purification from cells and tissue (Macherey-Nagel, Düren, Germany). The emptied crop was discarded to limit the carry-over of predator DNA in the gut content. At this point, samples were chosen in order to avoid specimens with visually empty crops. For all other specimens, gut content samples were stored at −20°C until further processing.

### 4 Library preparation and sequencing

DNA from gut contents samples was extracted using the epMotion 5075 workstation (Eppendorf, Hamburg, Germany) and the NucleoMag Tissue kit for purification from cells and tissue (Macherey-Nagel, Düren, Germany) following the manufacturer’s protocol. Initial gut content solution volume was 250 *μ*l, the final elution was performed in 100 *μ*l and the DNA extract was stored at −20 °C until PCR amplification.

A 313 bp fragment of the mitochondrial Cytochrome Oxidase I (COI) gene was amplified using the generic primer pair mlCOIintF (5’ - GGWACWGGWTGAACWGTWTAYCCYCC - 3’) and jgHCO2198 (5’ - TAIACYTCIGGRTGICCRAARAAYCA - 3’) (**Leray et al., 2013; Folmer et al., 1994**), correctly modified for high-throughput sequencing (HTS) on Illumina. PCR amplification reactions (25 *μ*l) contained the following: 2 *μ*l of template DNA, 1 *μ*l of each primer [10 *μ*M], 5 *μ*l of 5X GoTaq (Promega) reaction buffer, 1 *μ*l of MgCl_2_, 1 *μ*l of BSA, 0.5 *μ*l of dNTPs, 13.375 *μ*l of molecular-grade water and 0.125 *μ*l of GoTaq G2 Polymerase (Promega). PCR conditions were: 95 °C for 3 min, followed by 40 cycles of denaturation at 95 °C for 1 min, annealing at 48 °C for 1 min and elongation at 72 °C for 1 min 30 s, then a final elongation step was performed at 72 °C for 5 min. Amplification success was checked through a 1 % gel electrophoresis. Amplified amplicons were purified with a NGS clean up and size selection kit (Macherey-Nagel, Düren, Germany) following the manufacturer’s protocol. The final elution was performed in 40 *μ*L.

Later, the COI metabarcoding library was prepared by ligating Nextera XT indices through a ten cycle PCR (with the same conditions as for the initial PCR except for the annealing temperature which was 53°C for this second PCR). The concentration of the successfully ligated samples (checked on 1% agarose gel) was measured using a Qubit fluorometer (Life Technologies). Samples were then pooled in equimolar proportions (40 ng each), selected by size in a 1 % agarose gel and purified using the GeneJet Gel Extraction kit (Life Technologies), according to manufacturer’s protocol and the pools eluted in 30 *μ*l. Purified pools were combined into a 40 *μ*l final pool (4 nM). Sequencing runs were carried out on an Illumina Miseq using V2 chemistry (2 × 300 bp, 500 cycles) at the Sequencing Centre within the Biozentrum of the Ludwig-Maximilian-University in Munich (Germany). The raw dataset generated and analysed during the current study has been submitted to the NCBI Sequence Read Archive (SRA) under the BioProject PRJNA874542.

### 5 COI metabarcoding library filtering and taxonomic assignment

The FastQC software (https://www.bioinformatics.babraham.ac.uk/projects/fastqc/) was used to check the quality of the libraries (demultiplexed fastq files) on forward and reverse reads. Pair of primers were removed using *cutadapt* (**Martin, 2011**) and reads were merged with PEAR (**Zhang et al., 2014**), setting Phred score 30 as a threshold. The subsequent quality filtering (fastq_maxee = 1), dereplication, denoising, insertion and deletions (indels) and chimera removal were performed using the *vsearch v2.8.2* software (**Rognes et al., 2016**), which produced a fasta file containing Amplicon Single Variants (ASVs). These ASVs were clustered into Operational Taxonomic Units (OTUs) applying the centroid-based greedy clustering algorithm with a cut-off threshold of 97% (**Xiong** & **Zhan, 2018**) and an OTU table was mapped also using *vsearch v2.8.2* software (**Rognes et al., 2016**). Taxonomic classification of the sequences was performed against NCBI Genbank nr/nt using BLAST algorithm (**Johnson et al., 2008**) and R package *taxonomizr* (https://github.com/sherrillmix/taxonomizr/), in order to infer the species level classification when possible.

OTUs with a percentage of identity lower than 85% or a query coverage lower than 95% were considered as not assigned. Those with a percentage of identity between 85 and 93% were assigned to family, those between 93 and 97% to genus, and OTU with more than 97% percentage identity were assigned to species level.

### 6 Data analysis

The impact of trap type and sampling session on capture rate were analysed using generalized linear mixed effect models (GLMM) with Poisson distribution using the package *lme4* (**Bates** & **Maechler, 2013**). In this model, field ID was considered as a random effect. Likewise, the potential sex ratio differential between traps was assessed with a GLMM with binomial distribution with trap type and carabid species as fixed effects, and field ID and session as random effects.

To explore the trap type influence on the PCR success, we computed a binomial GLMM with trap type and species as explanatory variables. Field ID and session were considered as random effects. PCR success was defined by the presence/absence of visible amplified DNA band on the electrophoresis gel after 40 cycles of COI amplification and 10 cycles to attach the indices. A binomial GLMM was then performed on each species individually.

We estimated the completeness of the sampling by computing a rarefaction curve using R package *vegan* (**Oksanen et al., 2013**). The cumulative numbers of OTUs detected from individuals trapped in the classic pitfall traps and the dry pitfall traps for the two species were compared using a Kolmogorov-Smirnov test.

Next, the diet of carabids in oilseed rape crops was described using the Frequency of Occurrence (FOO) of prey OTU, which informs about the number of samples (counts) in which an OTU is present (**Deagle et al., 2019**). Only occurrences with at least two reads were considered. Relative Read Abundance (RRA), the proportion of reads of each OTU (**Deagle et al., 2019**), was not analysed in this study, since, most of the reads were predator reads while we were interested in prey. From the raw data, reads corresponding to carabids OTUs were thus discarded, since it was not possible to discriminate between the DNA of the predators themselves and that of other carabids which may have been predated. Contaminants OTUs (fungi, algae and taxa for which identity, prey or contaminant, cannot be decided) were also discarded. The FOO for prey OTUs was calculated using customized scripts using R package *dplyr* (**Wickham et al., 2021**).

To test the influence of the trap type on the sequencing results, we performed GLMM with Poisson distribution on the number of prey OTUs per carabid specimen, and a GLMM with binomial distribution on the sequencing success (as the presence/absence of at least one prey OTU per carabid specimen). Trap type and species were considered as fixed effects and field ID and the session as random effects.

Prey communities (beta diversity) of carabids caught between the two types of traps, were compared using a non-permutation multivariate analysis (PERMANOVA) with 999 iterations on the beta diversity using R package vegan (**Oksanen et al., 2013**).

All statistical analyses were performed with RStudio (R version 4.2.1).

## 7 Results

### 8 Trapping efficiency

A total of 611 carabid specimens from 14 carabid species were collected in oilseed rape fields, most of which (94.2 %) belonged to two species, *Nebria salina* (Fairmaire & Laboulbène, 1854) (n = 458) and *Calathus fuscipes* (Goeze, 1777) (n = 115) (**Table 1**). The other species were represented by a maximum of six individuals over the two sampling periods. Species richness was 0.84 (± 0.78 s.e.) in classic traps and 0.78 (± 0.72) in dry traps. These low diversity values do not allow to do a proper comparison of the two trap types in this respect.

**Table 1:** Carabid densities-activities and species richness recorded in classic (n=105) and dry (n=105) pitfall traps set up in 21 oilseed rape fields, with a focus on the dominant species Nebria salina and Calathus fuscipes

|  | Session 1 (5-9 Oct 2020) |  |  |  | Session 2 (25-29 Oct 2020) |  |  |  |
| --- | --- | --- | --- | --- | --- | --- | --- | --- |
|  | Density-activity |  |  | Species richness | Density-activity |  |  | Species richness |
|  | All carabids | *N. salina* | *P. cupreus* |  | All carabids | *N. salina* | *P. cupreus* |  |
| Classic traps | 151 | 98 | 43 | 8 | 154 | 134 | 14 | 6 |
| Dry traps | 103 | 51 | 39 | 10 | 203 | 175 | 19 | 7 |
| All traps | 254 | 149 | 82 | 12 | 357 | 309 | 33 | 9 |

The densities-activities were 1.78 (± 2.68) and 1.72 (± 2.00) in classic and dry pitfall traps respectively. We found a significant effect of the interaction between trap type and sampling session on the capture rate (t_60_ = −2.06, *p-value* = 0.044) but no effect of the trap type alone (t_60_ = 1.31, *p-value =* 0.195). Carabid density-activity was indeed higher in classic pitfall traps during the first sampling session whereas it was higher in the dry traps during the second sampling session (**Table 1**). Capture rate showed a strong bias in sex-ratio: of the 292 individuals used for the gut contents analysis, 215 were males and 77 were females (*N. salina*: 172, 55; *C. fuscipes*: 43, 22 ) but this bias was not related of the trap type (z = −1.04, *p-value* = 0.298).

### 9 PCR success

Out of 292 gut samples, 249 were successfully amplified for COI (187 *N. salina* and 62 *C. fuscipes*). PCR success rates were 92.3% in dry traps and 79.6% in classic ones. We found a significant effect of the species identity on PCR success (z = −2.07, *p-value* = 0.038). We found a higher PCR success with *C. fuscipes* specimens (95%) compared to *N. salina* specimens (82%). Performed separately on each species alone, PCR success was higher with *N. salina* individuals caught in dry pitfall traps (91.2%) than individuals trapped with pitfall traps (75.2%) (z = 2.139, *p-value* = 0.032). In contrast, there was no difference between trap types in *C. fuscipes* (classic 94.6%, dry 96.4%) (z = 0.34, *p-value* = 0.737).

### 10 Sequencing success

DNA was successfully sequenced from 97.2% of the samples. A total of 14,261,981 reads was obtained after clean-up. An average of 45,565 reads (±17,054 s.e.) were obtained per sample. The vast majority of those reads corresponded to carabids (95%). The remaining reads corresponded to prey (2,7 %), fungi (1.2%), algae (0.6%), other contaminants (0.2%) and non-assigned OTUs (0.3%).

The OTU accumulation curve suggested that the diet of the studied species was well estimated by our analysis with 82.4% of OTUs detected for *N. salina* and 89.9% for *C. fuscipes* (**Figure 2**). There were significant differences in the cumulative number of OTUs detected from individuals trapped with classic and dry pitfall traps for *Nebria salina* (KS, D = 0.35, *p-value*<0.0001) and *Calathus fuscipes* (D = 0.56, *p-value*<0.0001). Based on rarefaction curves, dry traps retrieved more OTUs than the classic traps for both *N. salina* and *C. fuscipes*.

**Figure 2:**
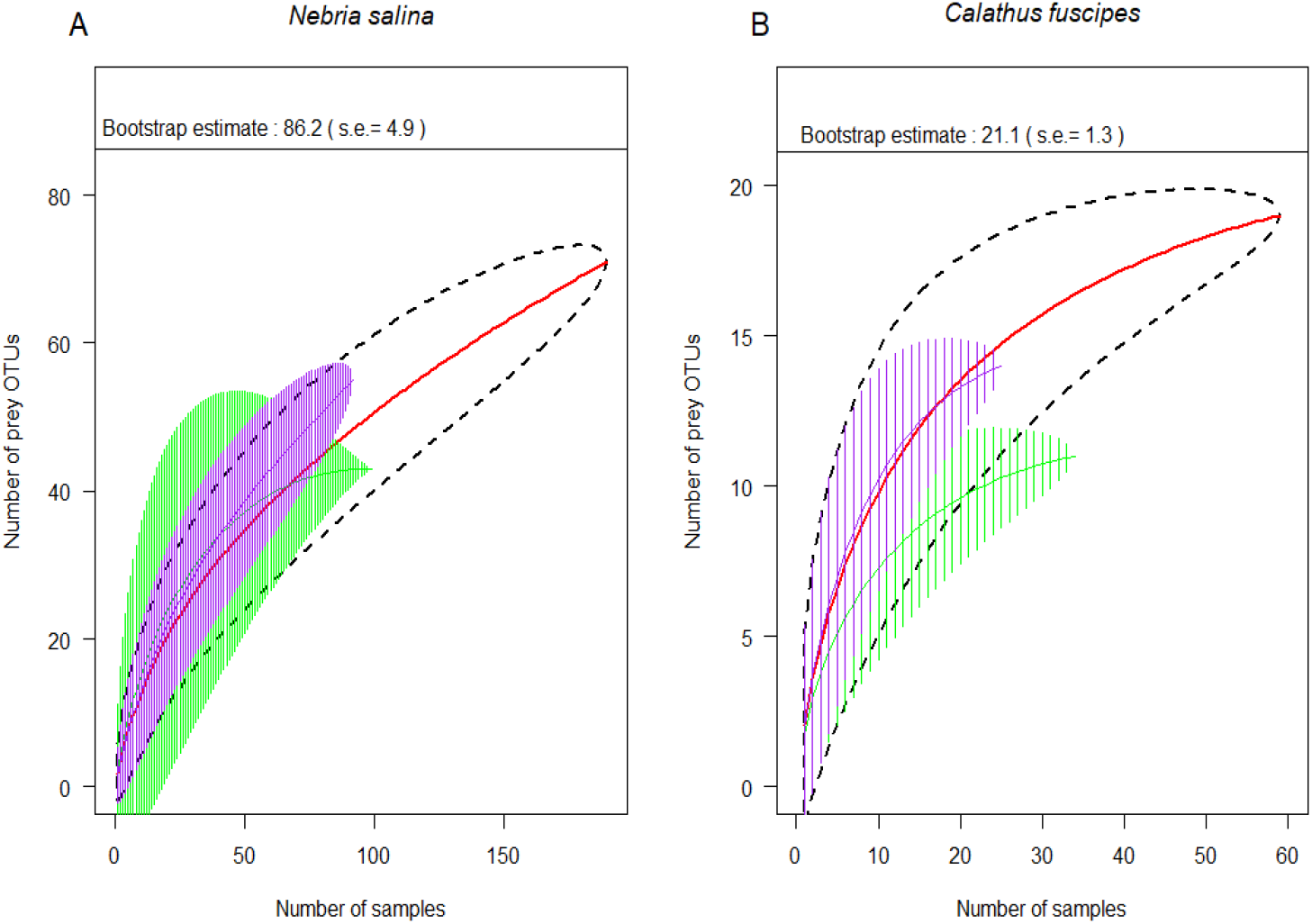
Detection of prey OTUs in the carabids gut contents. Cumulative curves of number of preys OTUs in relation to number of samples. (A) : Nebria salina, (B) : Calathus fuscipes. Cumulative curves are in red, the area delimited by the dashed lines corresponds to the 95% confidence interval. The horizontal solid lines represent the estimated total number of preys OTUs in the diet of carabids. The green and purple hatched areas correspond to cumulative curves obtained with classic and dry pitfall traps, respectively.

With regard to taxonomic assignation, 258 OTUs were assigned to contaminants. 95 OTUs were assigned to the Carabidae family, including 42 OTUs assigned to *Nebria salina* and *Nebria sp*., 33 to *Calathus fuscipes* and *Calathus sp*., 4 to Carabidae sp. and finally 16 OTUs were assigned to other carabid species (*Amara consularis, Anchomenus dorsalis, Harpalus rufipes, H. dimidiatus, Poecilus cupreus* and *Zabrus tenebrionides*). The remaining 142 OTUs matched prey DNA but 67 of them (representing only 237 reads) were discarded because they comprised only one read. Out of the 75 remaining prey OTUs, 68 were found in *N. salina* samples (44 in classic and 52 in dry pitfall traps) and 19 in *C. fuscipes* samples (11 in classic and 13 dry pitfall traps), with 12 OTUs in common between the two species. Based on their percentage of identity, 38 prey OTUs were assigned to species level, 15 to genus level and 22 to family level.

Prey OTU were found in 134 *N. salina* individuals out of 190, and 41 *C. fuscipes* individuals out of 59 (**Figure 3**). There were an average 1.75 prey OTUs per individuals in *N. salina* (classic: 1.84, dry: 1.65) and 1.36 in *C. fuscipes* (classic: 1.35, dry: 1.36). There was no influence of the type of traps on the sequencing success (z = 0.22, *p-value* = 0.83). Likewise, the number of prey OTUs per sample did not significantly differ between individuals trapped in the two type of traps for *N. salina* and *C. fuscipes* (z = 0.1, *p-value* = 0.62).

**Figure 3:**
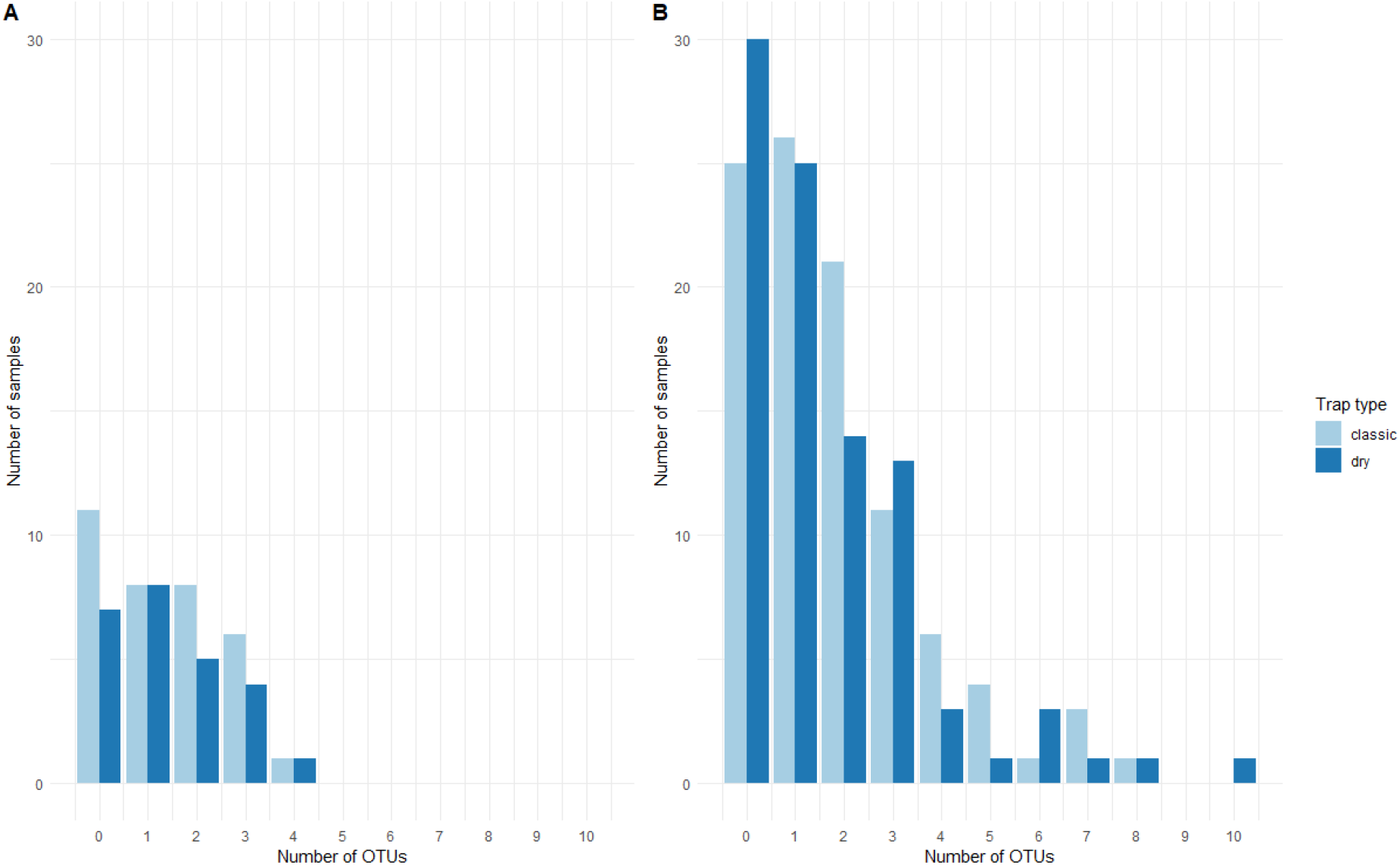
Distribution of carabid specimens collected in classic and dry pitfall traps, according to the number of prey OTU they contained. (A) Calathus fuscipes. (B) Nebria salina.

For *N. salina*, we found 15 invertebrate orders (14 in classic and 13 in dry pitfall traps) (**Figures 4a and 4b**). The frequency of occurrence was however different for the two types of traps. The main prey orders for samples trapped in classic pitfall traps were Entomobryomorpha (number of OTUs = 4; FOO = 43%), Opisthopora (5; 24%) and Diptera (8; 14%), whereas the main prey orders for the samples trapped in dry pitfall traps were Opisthopora (10; 27%), Diptera (18; 22%) and Entomobryomorpha (3; 22%).

**Figure 4:**
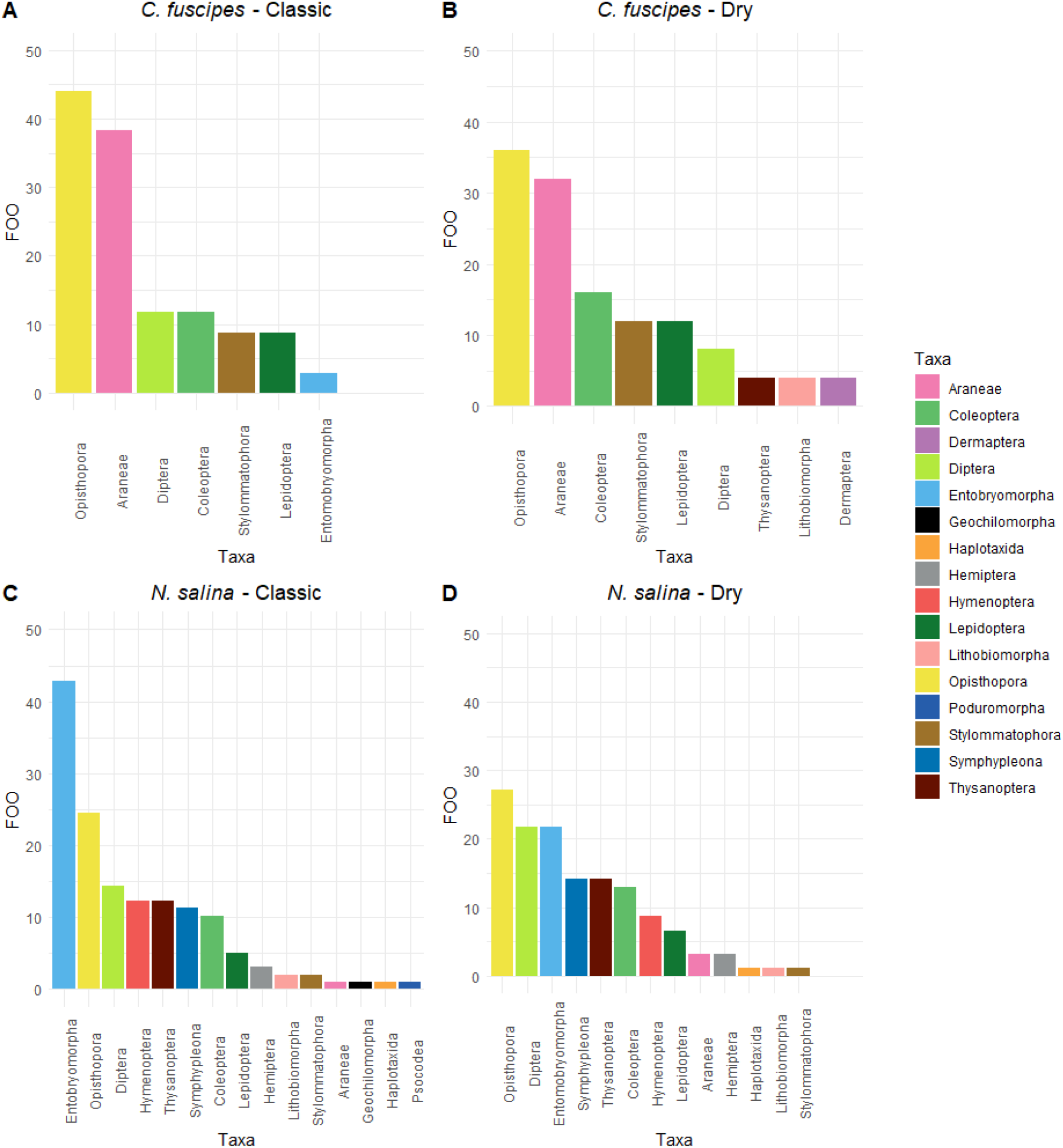
Frequencies of occurrence of invertebrate orders in the diet of Calathus fuscipes (A, B) and Nebria salina (C, D) captured in oilseed rape fields with classic (A, C) and dry pitfall traps (B, D). (A) : Diet of 24 individuals comprising 11 prey OTUs, (B) : Diet of 18 individuals comprising 13 prey OTUs, (C) : Diet of 74 individuals comprising 43 prey OTUs, (D) : Diet of 61 individuals comprising 55 prey OTUs.

For *C. fuscipes*, we found 10 invertebrate orders (9 in classic and 7 in dry pitfall traps) (**Figures 4c and 4d**). The relative contribution of each order in the diet of the specimens trapped in classic and in dry pitfall traps were similar. There was a major contribution by Opisthopora (especially by *Aporrectodea longa* Lumbricidae) present in 44% (classic) and 36% (dry) of the samples and by Araneae (only *Amaurobius erberi* Amaurobiidae) present in 38% and 32% of the samples.

When analysing the beta diversity, we found a difference in the prey OTUs composition for carabids trapped in classic and dry pitfall traps for *N. salina* (PERMANOVA, F = 1.46, R^2^ = 0.01, *p-value* = 0.047). We did not find such a difference for *C. fuscipes* (F = 1.06, R^2^ = 0.026, *p-value* = 0.352).

## 11 Discussion

Metabarcoding approaches are powerful tools for the characterization of food webs (**Pompanon et al., 2012**). The diet of a high number of specimens can be characterized in a unique sequencing run and in a cost-effective manner, without the need to estimate precisely the diet composition prior to performing the study (**De Sousa et al., 2019**). When applied on an arthropod community in the context of an agroecosystem, metabarcoding offers the potential to retrieve the trophic relationships between predators and prey (**Cuff et al., 2021; Sow et al., 2020**) or between parasitoids and hosts (**Sigut et al., 2017; Sow et al., 2019**), and subsequently to assess pest regulations and identify promising biocontrol agents. However, the use of such methods or technique is dependent on the quality of DNA, and therefore relies on its (hopefully low) degradation in samples (**Deagle et al., 2006**). Improving knowledge about insect trapping may help to lessen DNA degradation and, therefore, improve our understanding food networks within agroecosystems. When comparing the performance of dry versus «classic» pitfall traps, we found that the trap type did not have any effect on carabid species richness and prey OTUs sequencing. In both types of traps, the two mostly trapped species were *Nebria salina* and *Calathus fuscipes*, representing alone c.94 % of the carabids trapped. PCR success rate, however, was higher in dry pitfall traps compared to classic ones for *N. salina*, but no difference was observed for *C. fuscipes*. Moreover, dry traps allow a more effective prey detection.

Below, we discuss our results and propose methodological improvements for carabid trapping in agroecosystems.

### Trapping efficiency

We found that the density-activity of carabids was higher in classic pitfall traps during the first sampling session whereas it was higher in the dry traps during the second session. Indeed, the number of trapped carabids differed between the trapping session thanks to dry pitfall traps which captured twice as many individuals during the second session compared to the first while the classic traps captured about the same number. This difference comes especially from the increase in the density-activity of *Nebria salina* which does not compensate for the decrease in that *of Calathus fuscipes* between the two sampling sessions. This differential trapping can be due to the influence of the capture rate of two classic traps in one of the fields during the first sampling session (with 41 specimens trapped, i.e. 27% of the whole sampling by the 105 classic traps).

The number of carabid species collected per trap was very low probably because traps were only activated for one night. Thus testing whether the two types of traps are equally effective to capture carabid species was not possible using our data. However, the numbers obtained do not suggest any striking difference between the two types of traps in terms of species diversity, which contrasts with a previous study in which a higher species richness (but not higher abundance) of trapped specimens was recorded in killing pitfall traps compared to dry ones (**Weeks** & **McIntyre, 1997)**. The lack of differences in species richness between the two types of traps may be explained by the fact that we restrained the sampling to carabids and did not consider all arthropod species as did **Weeks and McIntyre (1997)**. Furthermore, the use of a killing agent may have impacted the activity-density and diversity observed in the latter study due to increased local humidity or attractive odours (**Woodcock, 2005**). Indeed, we took care by removing odours in our classic traps (odourless soap, beads and traps cleaned after each use). We noted a strongly biased sex ratio in both species and trap type towards males, which may result either from higher dispersal activity by males especially when searching for females for mating purpose, or from lower activity by females especially when satiated (**Szyszko et al., 2004;** **Wallin** & **Ekbom, 1994)**.

A major limitation of the dry trapping method is the potential predation inside the trap (**McKravy, 2018;** **Woodcock, 2005**). However, a very low number of studies have attempted to estimate this (**Mitchell, 1963; Roubinet *et al*., 2017**) and most authors conclude that it is an infrequent phenomenon (e.g., identified in four traps out of 100 by **Roubinet et *al*., 2017)**. Here we used clay beads in the dry pitfall to limit the encounters between captured prey. During our sampling sessions, the insects’ parts (e.g., wings, legs, antennae), which were sometimes found inside the trap seemed not to be the result of a predation, but rather the result of the perturbations which have occurred during the transportation (between the field and the laboratory) and the freezing of specimens prior to sorting.

### PCR success

PCR success was lower for *Nebria salina* collected with classic pitfall traps compared with those from dry pitfall traps. Upon dissection carabids’ crops were examined and we discarded samples where no leftover food was visible. Thus, empty crop cannot be the reason for the PCR failure. The killing agent, as used in classic traps, can have deleterious effects on the gut contents (**Schmidt et *al*., 2006;** **Szinwelski et *al*., 2012**). Ethanol has hydrolytic and oxidative effects on DNA (**Vink et al., 2005**). The killing agent can also act as an inhibitor leading to failed PCR during library preparation, such as Formaldehyde (**Gurdebeke** & **Maelfait, 2002**) or salt (**Davalieva** & **Efremov, 2010**), which was used in our classic traps. Specimens from classic traps could have swallowed salt water, which was then mixed with the crop contents. Moreover, pieces of soil and other fragments of vegetation, which are often found in traps, could bring other PCR inhibitors, such as humic acid (**Watson** & **Blackwell, 2000;** **Schrader at al., 2012**) and plant polysaccharides (**Demeke** & **Adams, 1992;** **Schrader et al., 2012**). *N. salina* is a smaller species than *C. fuscipes* and may have been more impacted by this issue. Even if prey DNA was totally degraded inside the crop, predator DNA is always collected with the gut contents and ensures PCR success with COI primers (especially because the carabid DNA is in good condition compared with the prey DNA). The fact that several PCR failed, means that even the carabid DNA could not be amplified, which supports the inhibitor hypothesis. We can also hypothesize that the difference between *N. salina* and *C. fuscipes* is linked to the diet of *N. salina*. Although we only amplified animal prey DNA, we could observe during dissections the presence of plant remains in the crops of *N. salina* but not in those of *C. fuscipes*. **Kamenova et al. (2018a)** also showed that *N. salina* frequently consume plant material. The presence of plant in the diet of *N.salina* is likely to enrich their crop contents with PCR inhibitors (**Dermeke** & **Adams, 1992; Schrader et al., 2012**).

### Sequencing success

Conversely to PCR, sequencing success (taken as the ability to retrieve at least one prey OTU) was not different between carabids caught in the two types of traps. Likewise, the number of prey OTUs was not different between individuals caught in the two trap types. When we analysed the beta diversity, we did not find a difference in prey OTUs composition between both types of traps for *C. fuscipes* but a slight difference for *N. salina*.

For *C. fuscipes*, the two major prey, *Amaraubius erberi* (a spider) and *Aporrectodea longa* (a worm), could be detected equally well and with the same rank order in the two trap types. For *N. salina*, the three main prey orders were always Opisthopora, Diptera and Entomobryomorpha, regardless of the type of trap. Only, the frequencies of occurrence differed. The more notable difference concerns Entomobryomorpha which were detected in 43% of individuals trapped with classic pitfall traps but only in 22% of individuals trapped with dry ones. This overrepresentation may result from the degradation of prey DNA during the carabid digestion, which continues in dry pitfall traps until the specimens are frozen. For medium sized DNA fragments (300-500 bp), the detectability half-life (T_50_ - an estimated time post-feeding for a 50% prey DNA detection probability) ranged around 30 hours post-feeding for carabids in **Waldner et *al*. (2013)** or around 40 hours post feeding in **Kamenova et al. (2018b**). Traps were activated for 15 hours at the most to limit prey digestion by the predators. However, it is likely that carabids had consumed one or more prey potentially hours before falling into the trap, which increased the risk that prey DNA had been digested and could not be detected. On the contrary, in classic traps, although digestion may continue for some time, degradation is probably slowed down. It seems that this problem was not acute in our case as accumulation curves and the prey OTUs numbers actually indicated that dry pitfall traps make it possible to reconstitute more effectively the prey community than classic pitfall traps.

As opposed to specimens trapped with concentrated ethanol as killing agent (**Szinwelski et *al*., 2012**), those collected in standard traps containing slow-killing agents (such as water or brine) can react with regurgitations or defecations (**King et al., 2008**) or by swallowing liquids. These reactions may cause cross contamination among the trapped specimens. **Athey et al. (2017**) on the contrary proved that a slug drowned inside ethanol for 24 hours cannot contaminate the gut contents of the carabid *Pterostichus melanarius*. Moreover, they could not detect any amplifiable slug DNA in the killing agent. Conversely, **Shokralla et al. (2010**) showed that even in absence of regurgitation or defecation, specimens start diffusing amplifiable DNA into ethanol after just 24 hours. It is not excluded that this killing agent can be a source of cross contamination, especially if many specimens are mixed in the liquid. This is probably the case with Collembola, which were abundantly trapped. Hence, the overrepresentation of Emtomobryomorpha DNA in specimens trapped in classic pitfall traps could be the consequence of cross contamination between Collembola and carabids soaking in the same liquid.

A small number of reads came from prey OTUs in comparison to reads coming from carabid OTUs. Because of the phylogenetic proximity between the predator and its prey, the PCR with universal primers leads to amplification of both the prey and the predator (**O’Rorke et al. 2011; Piñol et al., 2013**). Therefore, the competition with the predator DNA can prevent the amplification of prey DNA (**Piñol et al., 2013; Shehzad et al. 2012**). This situation is even more acute with a high degradation of prey DNA in the gut (**Cuff et al., 2022**). On the contrary, the predator DNA is generally in good condition, and despite the precautions taken during the dissection, it can remain in high quantity inside the sample as compared to prey DNA. Therefore, more predator DNA is amplified during the PCR compared to prey DNA. One solution could be to develop blocking primers to prevent the amplification of predator DNA (**Vestheim** & **Jarman, 2008**). However, developing efficient blocking primers is not easy, especially for carabids **(Kamenova, 2013**).

An alternative solution is to lower the detection threshold to avoid rejecting any prey DNA occurrence and retrieve as much dietary information as possible. When following this strategy, prey OTUs are represented by low numbers of reads, and it can be difficult to distinguish data output from potential contamination (**Drake et al., 2022**). That is why special attention has been taken to prevent contaminations between samples or with the laboratory equipment during the sample preparation for sequencing. In our analysis, no prey DNA was detected from the blanks, suggesting that the contamination risks were properly mitigated.

### 12 Prey spectrum of N. salina and C. fuscipes in Autumn

These two species are well known to be autumn breeders (**Holland, 2002**), active at night (**Kegel, 1990; Bargmann et al., 2016)** and considered as generalist predators (**Luff, 2002; Bargmann et al., 2016**). Their diet is not well known, and therefore our study provides key information about their prey spectrum in autumn. *Nebria salina* is known to consume plants, Diptera and Araneae in wheat and oilseed rape fields (**Kamenova et al., 2018a)**. The species is morphologically very similar to *N. brevicollis* (Fabricius, 1792), whose diet is more often studied and seems mainly composed of Collembola and mites, along with other soil-living Arthropods as accessory food (**Penney, 1966;** **Toft** & **Bilde, 2002**). In oilseed rape, *N. salina* showed here a clear preference for decomposers, Collembola, mainly Isotomidae (Entomobryomorpha) and Sminthuridae (Symphypleona), and earthworms (Opisthopora Lumbricidae). Collembola is one of the most abundant groups of Arthropods in agrosystems and recognized as a major prey group for carabids (**Mitchell, 1963;** Hengeveld, 1980; **Pollet** & **Desender,1987**). These organisms are generally considered as a low-quality food for adult carabids (**Bilde et al., 2001**) and are not easy prey (due to their escaping abilities), which has led certain predatory species to develop specific morphological adaptations to hunt Collembola (**Baulechner et al., 2021**) or to consume dead springtails (**Bilde et al., 2001**). *Nebria spp*. are not considered as Collembola specialists [1] even if the number of individuals having consumed springtails seems to indicate a certain hunting efficiency for this prey, or at least a frequent scavenging. Preyed Diptera belonged mostly to the families Sciaridae and Chironomidae, which are among the main Diptera taxa found in agricultural landscapes (**Delettre** & **Lagerl, 1992;** **Frouz, 1997)**. *Episyrphus balteatus* (Syrphidae) is also a Diptera prey often encountered. Although these results suggest a probably weak role of *N. salina* as pest regulator, and even a potential negative effect on decomposers, the predation on non-pest prey could maintain carabid populations within the field during the pest dearth periods, thereby allowing predation on pests during the following spring (**Harwood** & **Obrichi, 2005;** **King et al., 2010**). Thus, it is necessary to assess the diet of *N. salina* in spring, when it is the second most abundant species in oilseed rape in the ZAPVS, to know whether *N. salina* turns into a pest predator.

*Calathus fuscipes* on the other hand can predate insect pupae, and fruits (**Larochelle, 1990**). In our study, the species seems to be more oligophagous than *N. salina*. Two main prey species are predated, the generalist predator *Amaurobius erberi* (Keyserling, 1863) (Araneae Amaurobiidae) and the anecic worm *Aporrectodea longa* (Ude, 1885) (Opisthopora, Lumbricidae). As for *N. salina*, this prey spectrum could lead to reject *C. fuscipes* as potential biocontrol agent. However, the relatively weak number of individuals analysed calls for caution when concluding on these predations on one generalist predator and one decomposer. Moreover, *C. fuscipes* emerges at the end of the spring and it may have a different diet during the summer.

In conclusion, our data showed that the trapping efficiency is under the influence of the trap type and the sampling session in regard to the density-activity. PCR success rate was higher only for *Nebria salina* specimens caught in dry pitfall traps compared to individuals caught in classic ones probably because of the combined effects of inhibitors present in the classic pitfall traps and in the digestive tract of *N. salina* due to its diet containing more plants.

In addition, the ability to retrieve prey OTUs was higher for dry pitfall traps. As suggested by the accumulation curves and the beta diversity for *N. salina*, the prey OTUs composition is higher for specimens caught in dry pitfall traps.

Although frequencies of occurrence varied between the two trapping methods, the list of prey taxa detected were almost identical for individuals of the same species whether they were caught in classic or in dry pitfall traps. Since Collembola were trapped in larger quantities, we can also hypothesize that the overrepresentation of Entomobryomorpha in the diet of carabids trapped in classic pitfall traps could be the result of contamination through the killing agent.

As compared to classic ones, dry pitfall traps present the added advantage of being more focused on the target taxa and less invasive for invertebrate communities. Furthermore, the capture of live specimens presents an alternative to collect regurgitates and release the insects (**Waldner** & **Traugott, 2012**). These advantages make dry pitfalls a preferred choice, especially when repeated or intensive sampling is required. However, in this study, we cannot totally reject the hypothesis of internal trap predation. Therefore, future experiments should aim at measuring this phenomenon in dry pitfall traps.

## 13 Author contributions

Y.G. and S.B. designed research; V.B. and S.G. provided logistic support for the fieldwork.Y.G. collected the samples and performed the laboratory work. M.Q. performed the bioinformatic analyses. Y.G. and M.Q. conducted the statistical analyses. Y.G. and S.B. wrote the manuscript. All authors contributed substantially and critically to revisions.

## Acknowledgments

This work was supported by the ANR funded project IMAgHO (ANR-18-CE32-0002-01).

